## Supplementary material for "Critical Role for Isoprenoids in Apicoplast Biogenesis by Malaria Parasites": Figure Supplements

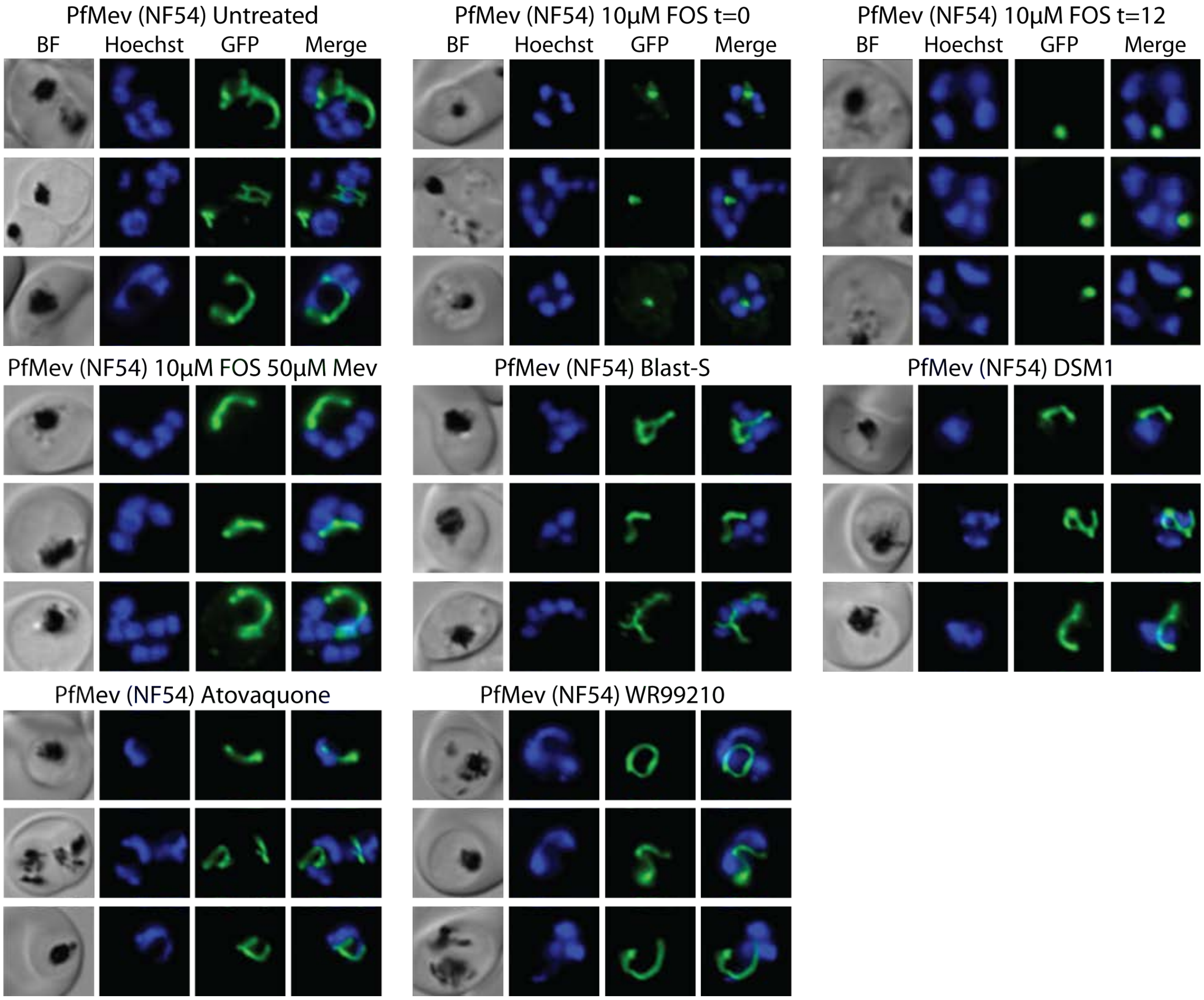


**Figure 1- figure supplement 1.** Additional epifluorescence images of PfMev $ACP_L$-GFP NF54 parasites treated with FOS and other drugs at the indicated concentrations. Parasites were synchronized to the ring stage with 5% D-sorbitol and incubated with the indicated treatments for 38 hours prior to live-cell imaging.

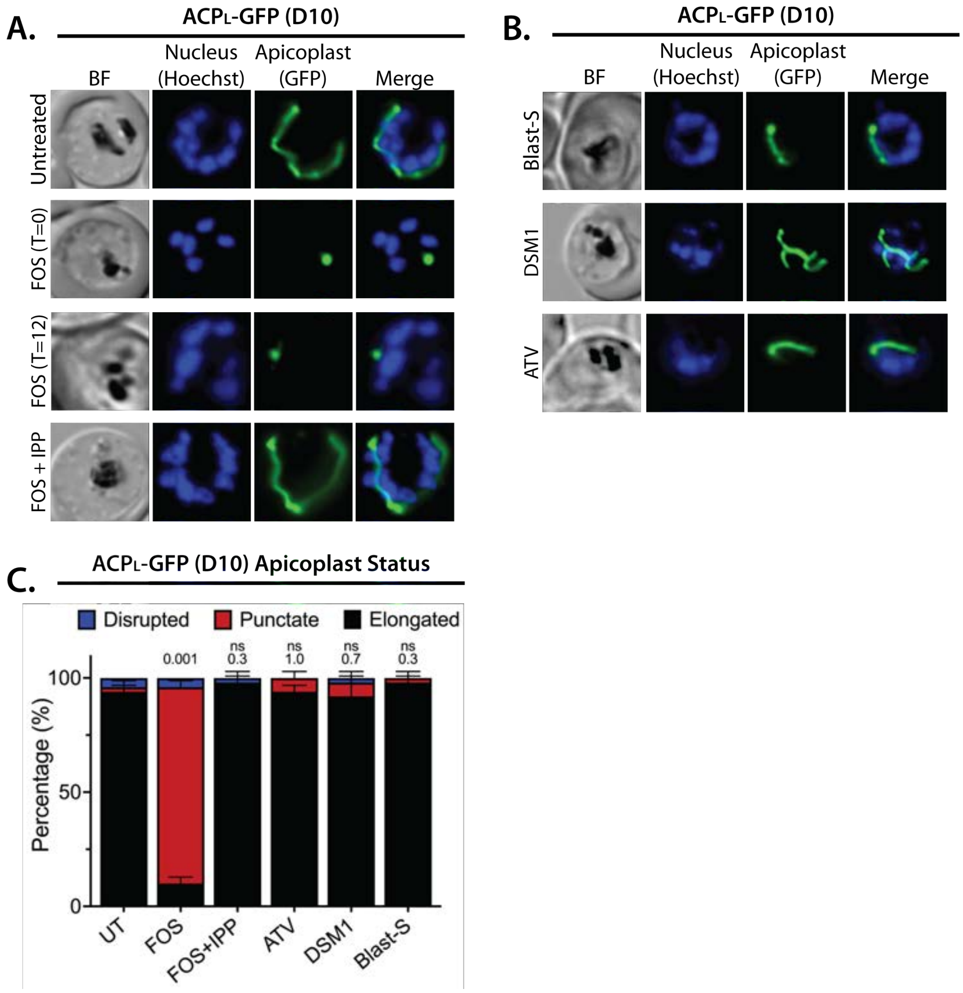


**Figure 1- figure supplement 2.** Epifluorescence images and analysis of D10 ACPL-GFP parasites treated with FOS and other drugs. Bright field (BF) and fluorescent microscopy images of live ACPL-GFP D10 parasites that were **(A)** untreated or treated with 10 µM fosmidomycin (FOS) in the absence or presence of 200 µM IPP, or **(B)** treated with 6 µM blasticidin-S (Blast-S), 2 µM DSM1, or 100 nM atovaquone (ATV). **(C)** Statistical analysis of apicoplast morphology for 50 total parasites imaged for each condition in panels A and B from two independent experiments. Apicoplast morphologies were scored as punctate (focal), elongated, or disrupted (dispersed); counted; and plotted by histogram as the fractional population with the indicated morphology. Error bars represent standard deviations from replicate experiments. Two-tailed unpaired t-test analysis was used to determine the significance of observed population differences compared to untreated (UT) parasites (P values given above each condition, ns = not significant). In all experiments, synchronized ring-stage parasites were incubated with the indicated treatments for 36 hours prior to live-cell imaging. Parasite nuclei were visualized using 1 µg/ml Hoechst 33342. The parasite apicoplast was visualized using the ACPL-GFP encoded by the D10 line.

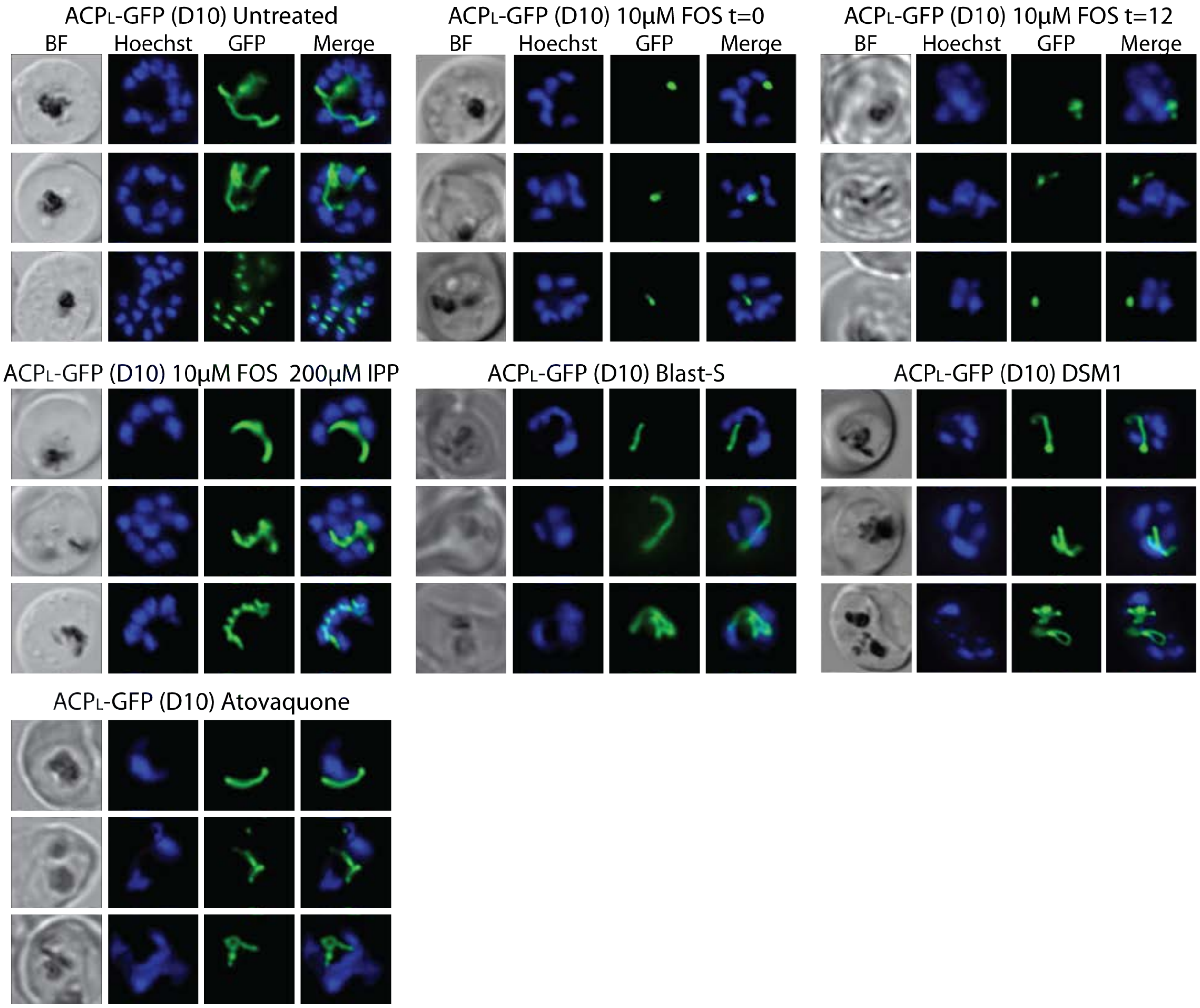


**Figure 1- figure supplement 3.** Additional epifluorescence images of D10 parasites treated with FOS and other drugs.

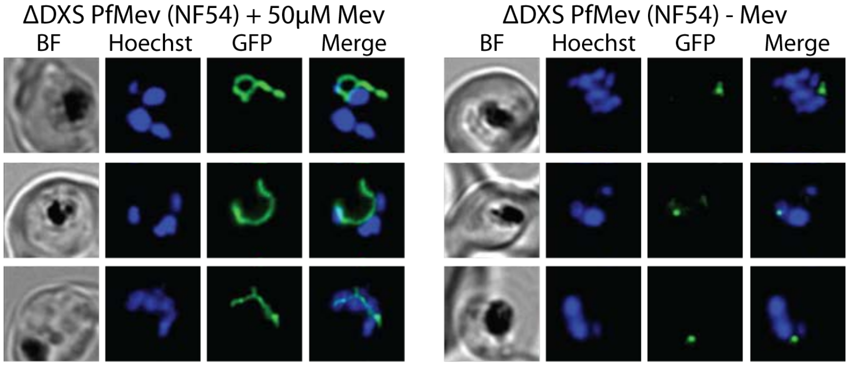


**Figure 1- figure supplement 4.** Additional epifluorescence images of ΔDXS PfMev parasites that were synchronized to ring stage with 5% D-sorbitol and incubated for 36 hours ±Mev hours prior to live-cell imaging. Parasite nuclei were visualized using 1 µg/ml Hoechst 33342. The parasite apicoplast was visualized using the $ACP_L$-GFP encoded by the PfMev line.

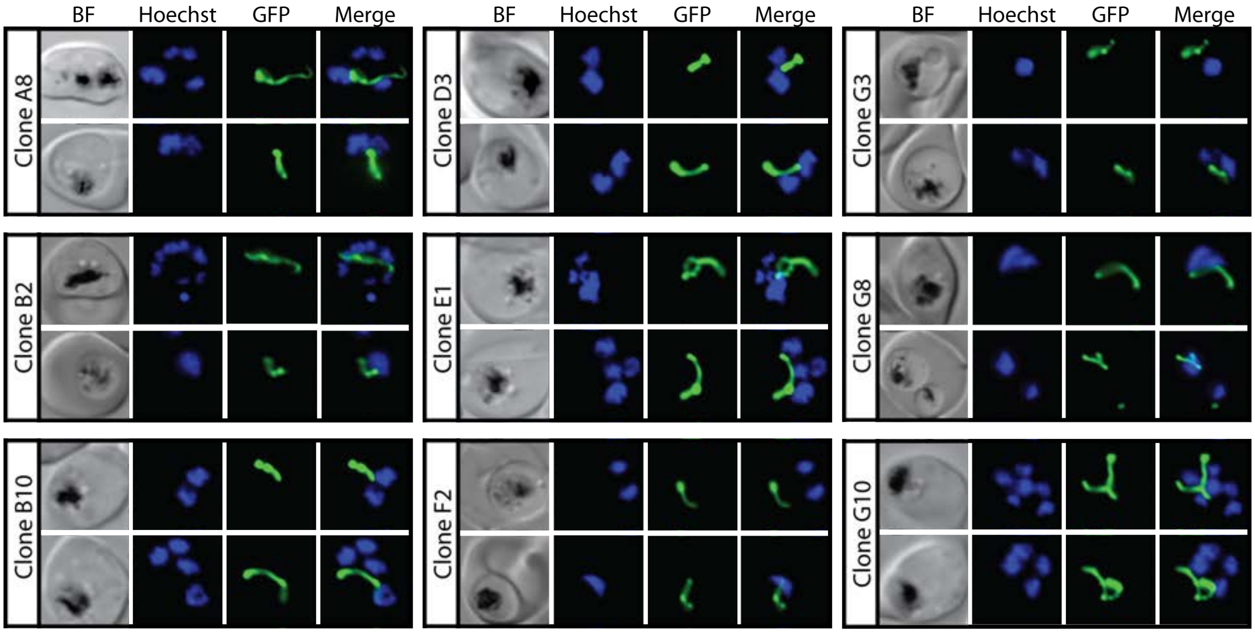


**Figure 2- figure supplement 1.** Epifluorescence microscopy images of clonal parasites isolated after FOS treatment and rescue by mevalonate addition at 0 hours after synchronization.

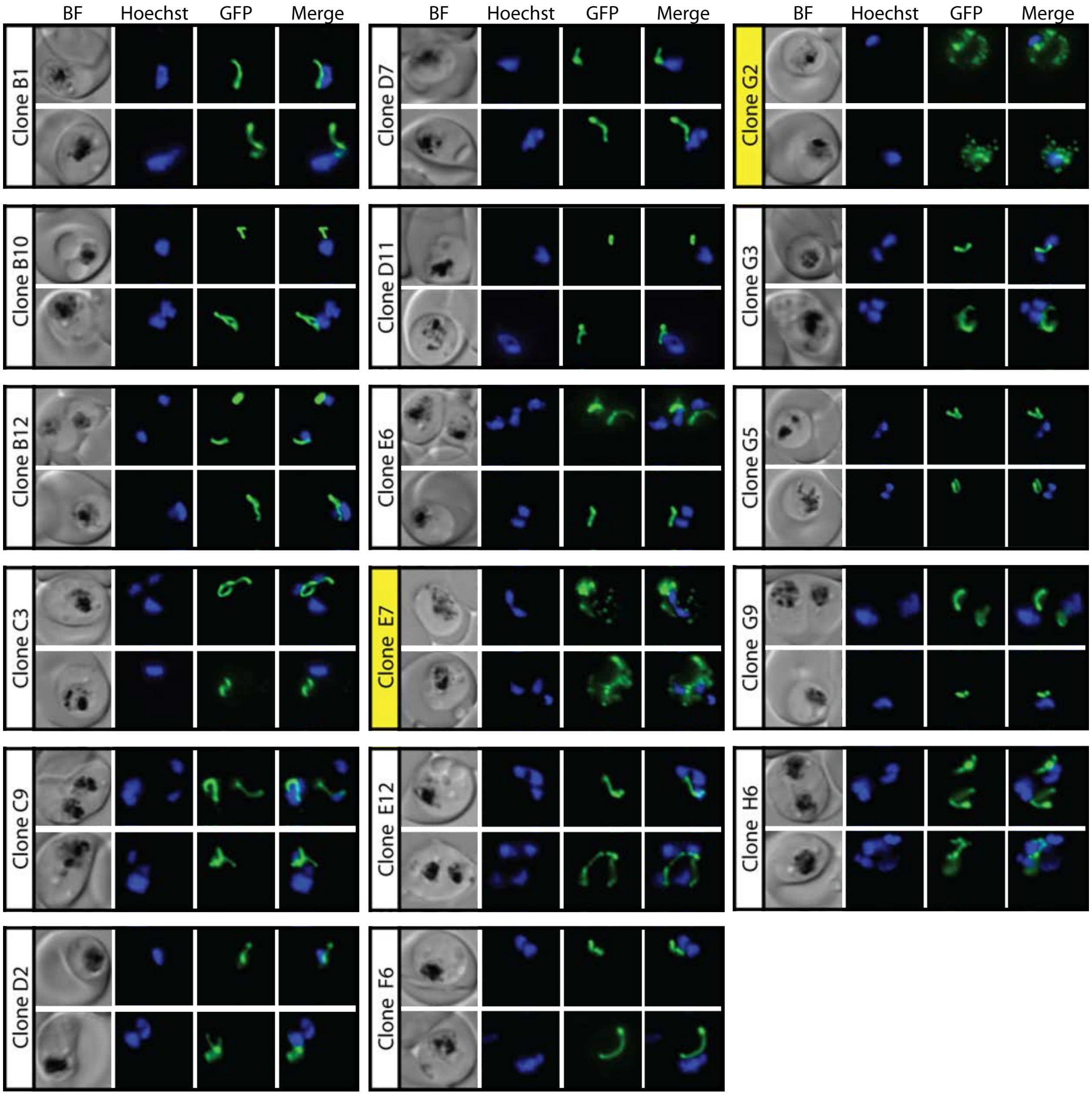


**Figure 2- figure supplement 2.** Epifluorescence microscopy images of clonal parasites isolated after FOS treatment and rescue by mevalonate addition at 30 hours after synchronization. Clone headers for parasites with a disrupted apicoplast are yellow.

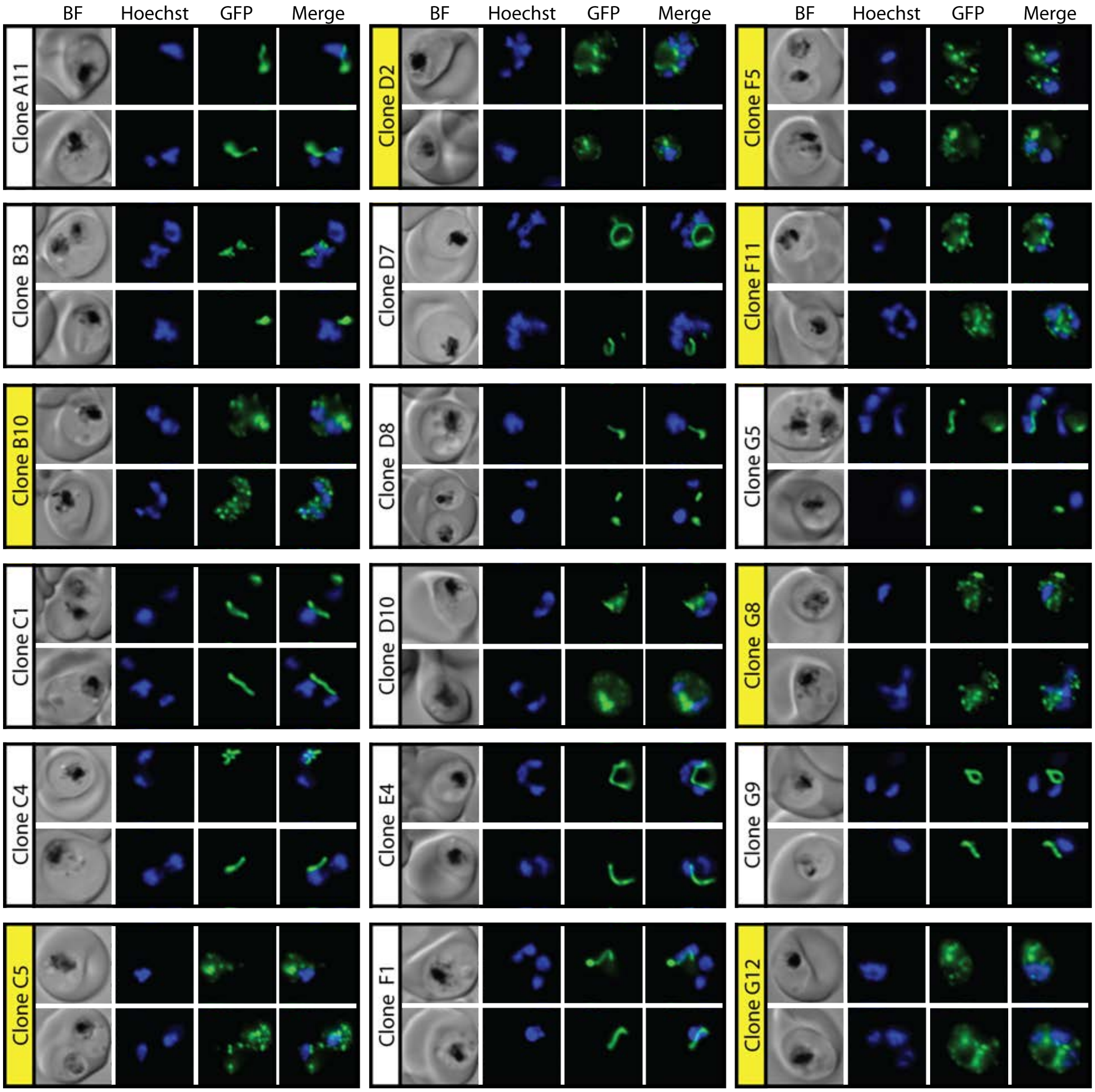


**Figure 2- figure supplement 3.** Epifluorescence microscopy images of clonal parasites isolated after FOS treatment and rescue by mevalonate addition at 34 hours after synchronization. Clone headers for parasites with a disrupted apicoplast are yellow.

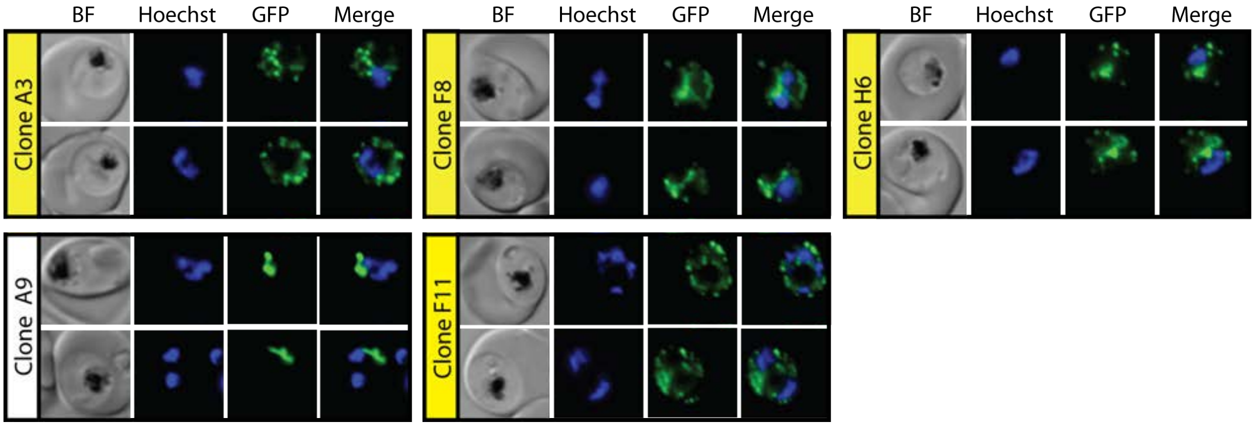


**Figure 2- supplement 4.** Epifluorescence microscopy images of clonal parasites isolated after FOS treatment and rescue by mevalonate addition at 38 hours after synchronization. Clone headers for parasites with a disrupted apicoplast are yellow.

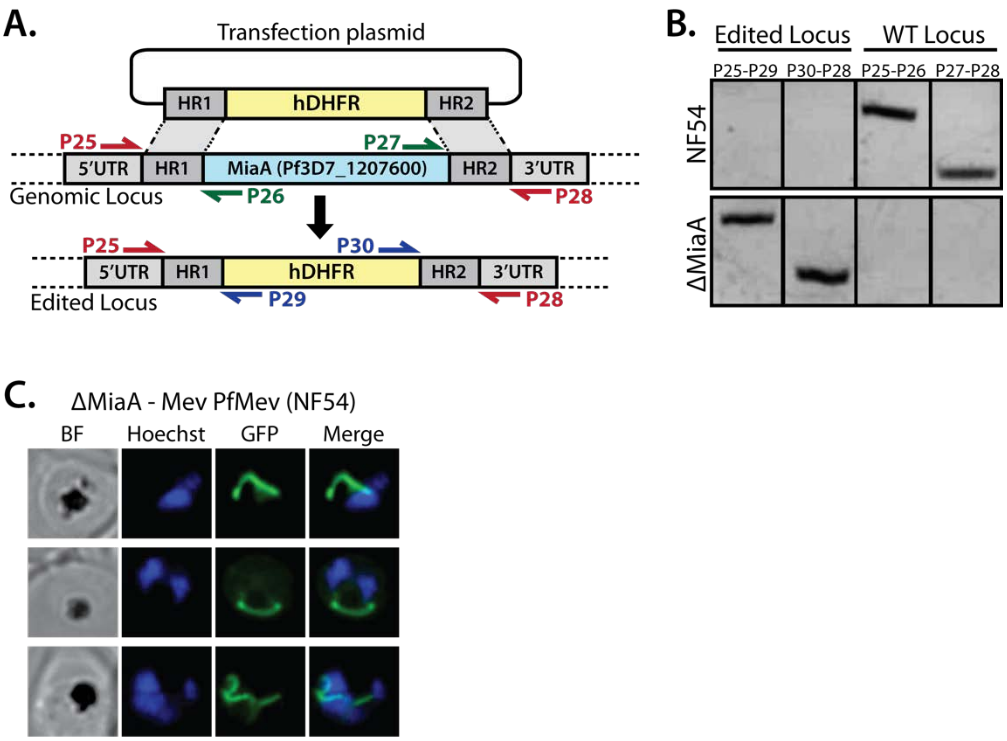


**Figure 3- supplement 1.** PCR genotyping of PfMev ΔMiaA parasites and additional epifluorescence images of apicoplast morphology. **(A)** Schematic depiction of the MiaA gene-disruption strategy using CRISPR/Cas9 and positive selection with human DHFR. Colored arrows depict PCR primer pairs used in panel B to test for retention or disruption of the MiaA gene. **(B)** Genomic PCR analysis of parental NF54 PfMev parasites and polyclonal transfected parasite progeny confirmed successful disruption of the MiaA gene. **(C)** Additional images of ΔMiaA parasites.

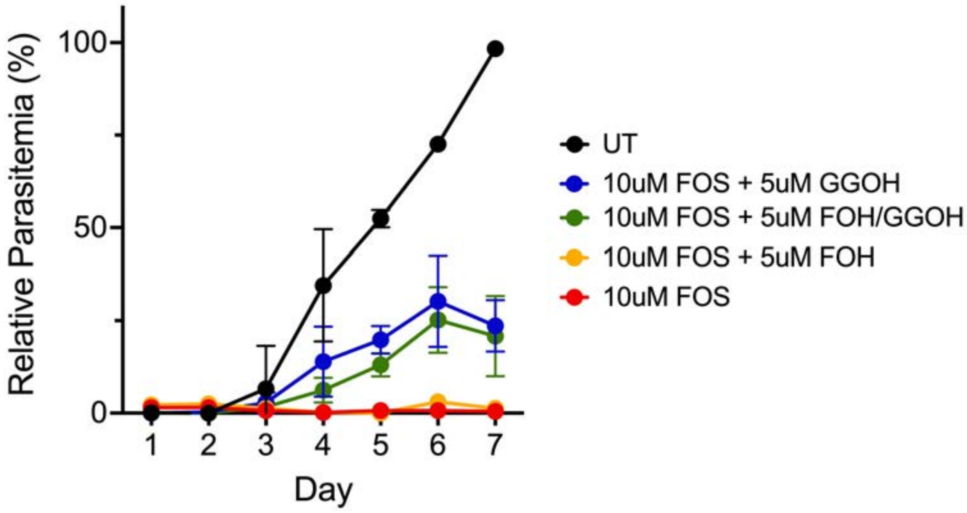


**Figure 4- figure supplement 1.** 5 µM GGOH but not FOH partially rescues parasite growth from inhibition by 10 µM FOS in continuous-growth assays with PfMev parasites. Data points are the average ±SD of two biological replicates. Parasites were synchronized to ring stage with 5% D-sorbitol and incubated with the indicated treatments. Daily parasitemia values were determined by flow cytometry.

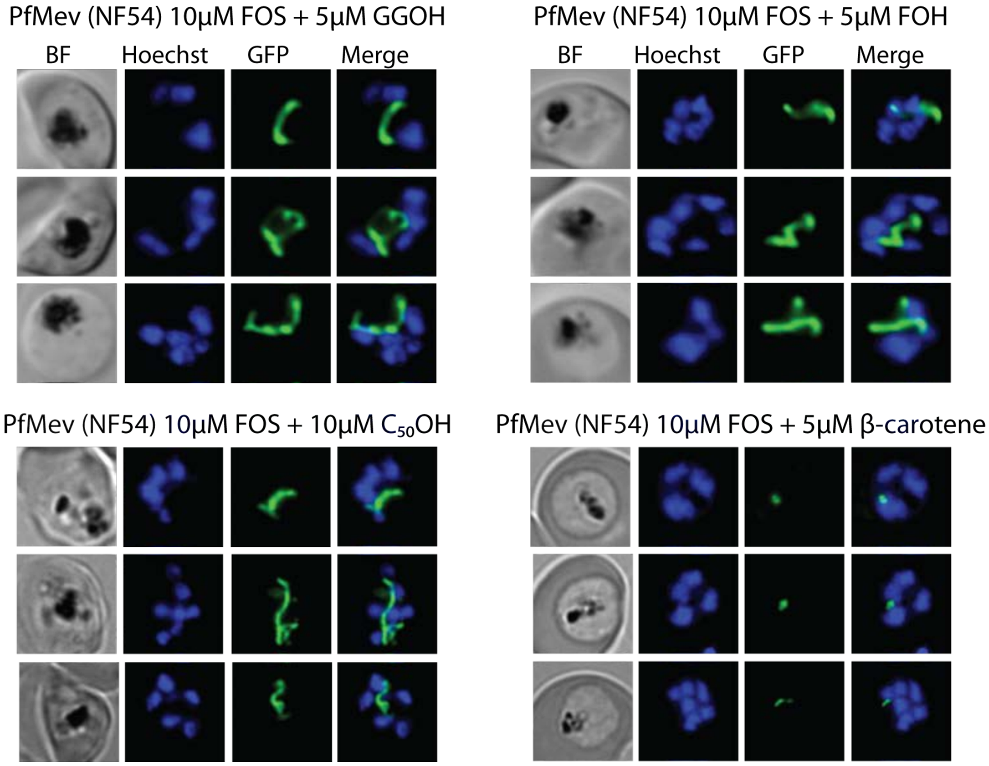


**Figure 4- figure supplement 2.** Additional epifluorescence microscopy images of PfMev parasites treated with FOS and FOH, GGOH, $C_{50}$-OH, or β-carotene. Synchronized ring-stage parasites were incubated with the indicated treatments for 36 hours and imaged by brightfield (BF) or fluorescence microscopy, with visualization of parasite nuclei by Hoechst staining and the apicoplast by $ACP_L$-GFP signal.

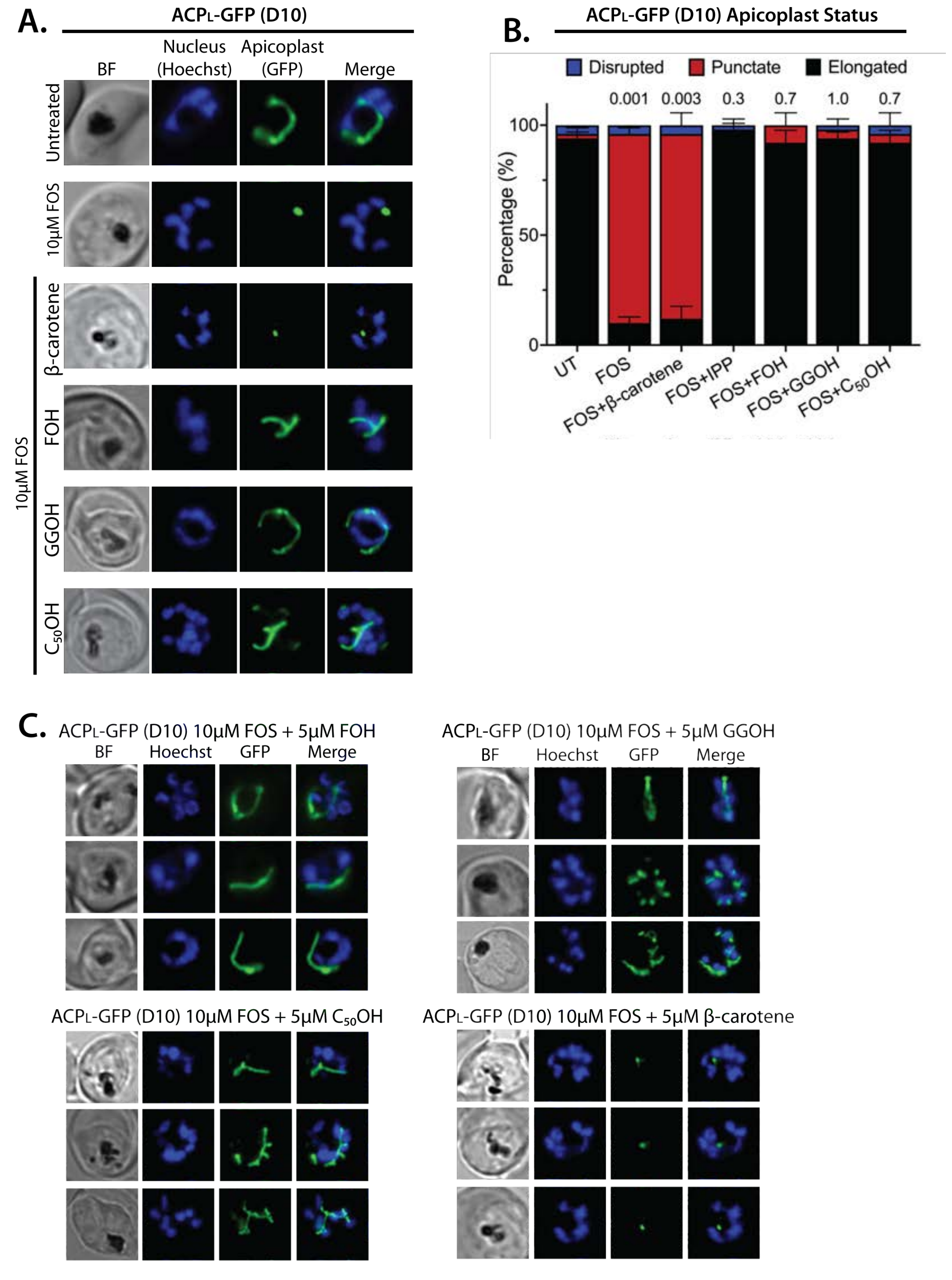


**Figure 4- figure supplement 3.** Epifluorescence microscopy images of D10 $ACP_L$-GFP parasites treated with FOS and FOH, GGOH, $C_{50}$-OH, or β-carotene. **(A)** 5 μM farnesol (FOH), geranylgeraniol (GGOH), or decaprenol ($C_{50}$-OH) but not β-carotene, rescues apicoplast biogenesis from inhibition by 10 μM FOS. **(B)** Statistical analysis of apicoplast status in D10 parasites treated with 10 μM FOS (P values are for comparison to untreated, UT). **(C)** Additional images of D10 parasites treated with FOS and FOH, GGOH, $C_{50}$OH, or β-carotene.

```
TR|O96130|O96130_PLAF7 MVHLSKRNNIKSFLNYCKAKYLNPLLINKNEDIIKETSILKNNNLYSRKESNVFIEILKS 60
SP|P08836|FPPS_CHICK   ------------------------------------------------------------
TR|Q8II79|Q8II79_PLAF7 ------------------------------------------------------------


TR|O96130|O96130_PLAF7 SFIKFRGQKINEEINNHNNIINNSSHNNNHNIYHDTNKKKKQQYEEKHNVFHTENMHKEV 120
SP|P08836|FPPS_CHICK   -MHKFTGVN---------------------------------------AKFQ-----QPA 15
TR|Q8II79|Q8II79_PLAF7 ------------------------------------------------------------


TR|O96130|O96130_PLAF7 LLCMDVLQYEEDKVNRELHLLHSYFNKE--------RTNIDPYTLCESKIKNIDEYIYNI 172
SP|P08836|FPPS_CHICK   LRNL-----SPVVVEREREEFVGFFPQIVR---DLTEDGIGHPEVGD------------- 54
TR|Q8II79|Q8II79_PLAF7 ---M-----ENEQNNQDSENGLDYFRSMYDRYRDVFINHINDYVLEDD--------IKII 44
                          :     .    ::: .   .:* .            *.   : :

TR|O96130|O96130_PLAF7 IKTNYKNIDEFITYIYLYKGKRFRVILSILLKNILHHIDNVSKIKTNFKNRNIQRKFFKS 232
SP|P08836|FPPS_CHICK   AVARLKEV---LQYN-APGGKCNRGLTVVAA----------------------------- 81
TR|Q8II79|Q8II79_PLAF7 ISKYYKLL---FDYN-CLGGKNNRGILVILI----------------------------- 71
                            * :   : *     **  * :  :

TR|O96130|O96130_PLAF7  NKLTSNYLSNKLKLYNLKITQKKNICEKTVLDNQCKIIAASEIIHMGSLLHDDVIDDSNK 292
SP|P08836|FPPS_CHICK    --------------YRE--LSGPGQKDAESLRCALAVGWCIELFQAFFLVADDIMDQSLT 125
TR|Q8II79|Q8II79_PLAF7  --------------YEY--VKNRDI-NCNEWEKVACIAWCIEILQASFLVADDIMDKGET 114
                                      *.    .  .  :         :  . *:::   *: **::*.. .

TR|O96130|O96130_PLAF7 RRGVIALHKKFGNKISILSGD-YLLARASSIFAGTGSPKICRSFSYVVE----------S 341
SP|P08836|FPPS_CHICK   RRGQLCWYKKEGVGL-DAINDSFLLESS--VYRV--LKKYCRQRPYYVHLLELFLQTAYQ 180
TR|Q8II79|Q8II79_PLAF7 RRNKHCWYLLKDVEIKNAVNDVFLLYNA--IYKL--LDVYLRNDNCYLDLITSFREATLK 170
                       **.  . :   .  :    .* :**  :  ::         *.    :.          .

TR|O96130|O96130_PLAF7 LIKGEFLQRNLK------------FNNV------EEALKMYLIKSY-----HKTA--SLF 376
SP|P08836|FPPS_CHICK   TELGQMLDLITA-------------------PVSKVDLSHFSEERYKAIVKYKTAFYSFY 221
TR|Q8II79|Q8II79_PLAF7 TIVGQHLDTNIFSDKYSHIDKDIDVNNINISQENKININMLNFKVYQNIIIHKTAYYSFF 230
                          *: *:                          :  :.    : *     :***  *::

TR|O96130|O96130_PLAF7 SHLFACIAILSFKND-TIIQLCFNLGLHIGMAFQLYDDYLDYKIDDNTNKPILNDLKNNI 435
SP|P08836|FPPS_CHICK   LPVAAAMYMVGIDSKEE-HENAKAILLEMGEYFQIQDDYLDCFGDPALTGKVGTDIQDNK 280
TR|Q8II79|Q8II79_PLAF7 LPIVCGMQMGGISLDNLLYKKVENIAILMGEYFQVHDDYIDTFGDSKKTGKVGSDIQNNK 290
                         : . : : .:. .    :    : : :*  **: ***:*   *   .  : .*:::*

TR|O96130|O96130_PLAF7 KTAPLLFSYNYNPQVILQLINKNSYTNNDIENILYY---------------IQHSNSMKK 480
SP|P08836|FPPS_CHICK   CSWLVVQCLQRVTPEQRQLLEDN-YGRKEPEKVAKVKELYEAVGMRAAFQQYEESSYRRL 339
TR|Q8II79|Q8II79_PLAF7 LTWPLIKAFELCSQPEKEDIIRN-YGKDNVTCIKFINDIYEHYNIRDHYVEYEKKQKMKI 349
                        :  :: . :       : :  * * ..:   :                   :...   :

TR|O96130|O96130_PLAF7 NELCSLLHIKKASDILYSLISHCNKPSTNKNNTKHDDIKQSSEALINLILNVLSRNVK-- 538
SP|P08836|FPPS_CHICK   QELIEKHSNR----------------------------------LPKEIFLGLAQKIYKR 365
TR|Q8II79|Q8II79_PLAF7 LEAINQLHHE--------------------------GIEYVLKYVMDILFTGA------- 376
                        *  .    .                                  : . ::

TR|O96130|O96130_PLAF7 --
SP|P08836|FPPS_CHICK   QK 367
TR|Q8II79|Q8II79_PLAF7 --
```

**Figure 5- figure supplement 1.** Full sequence alignment of PF3D7_0202700, PF3D7_1128400, and avian FPPS (Uniprot P08836).

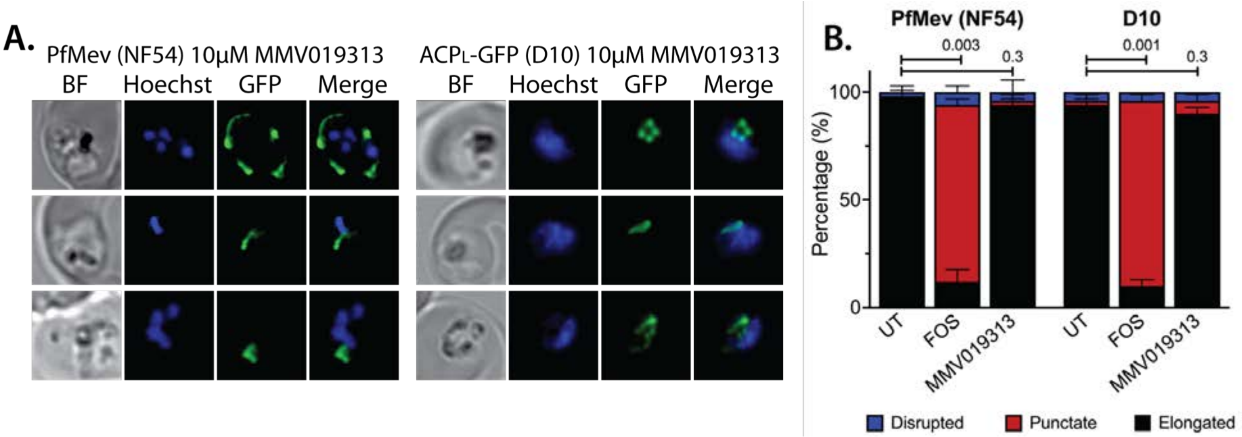


**Figure 5- figure supplement 2.** Epifluorescence microscopy images and statistical analysis of PfMev and D10 parasites treated with 10 µM MMV091313. **(A)** Synchronized ring-stage parasites were incubated with 10 µM MMV091313 for 36 hours and imaged by brightfield (BF) or fluorescence microscopy, with visualization of parasite nuclei by Hoechst staining and the apicoplast by $ACP_L$-GFP signal. **(B)** Statistical analysis of apicoplast morphology for 50 total parasites imaged for each condition in panel A from two independent experiments. Apicoplast morphologies were scored as punctate (focal), elongated, or disrupted (dispersed); counted; and plotted by histogram as the fractional population with the indicated morphology. Error bars represent standard deviations from replicate experiments. Two-tailed unpaired t-test analysis was used to determine the significance of observed population differences compared to untreated (UT) parasites (P values given above each condition).

Top 10 NCBI Blast results for PF3D7_02020700*,**

| Description | Species | % Ident | Accession |
|---|---|---|---|
| Geranylgeranyl pyrophosphate synthase | Haematococcus lacustris | 30.17% | KFY27494.1 |
| Decaprenyl-diphosphate synthase subunit 1 | Fistulifera solaris | 29.52% | APX64486.1 |
| Decaprenyl-diphosphate synthase subunit 1 | Strongylocentrotus purpuratus | 29.38% | XP_030829069.1 |
| Trans-prenyltransferase 4 | Haslea ostrearia | 29.94% | AYV97144.1 |
| Solanesyl diphosphate synthase 3 isoform X2 | Brassica napus | 30.54% | XP_006410610.1 |
| Solanesyl diphosphate synthase 1 isoform X1 | Asparagus officinalis | 28.51% | XP_020277177.1 |
| Solanesyl diphosphate synthase 3 isoform X2 | Arabidopsis lyrata | 30.24% | XP_02883166.1 |
| Geranyl diphosphate synthase 1 | Arabidopsis thaliana | 29.94% | NP_850234.1 |
| Solanesyl diphosphate synthase 3 | Capsella rubella | 30.65% | XP_023639436.1 |
| Solanesyl diphosphate synthase 1 | Physcomitrella patens | 28.10% | XP_024 |

*excluding Apicomplexan (taxid:5794)
**excluding predicted and hypothetical proteins

Top 10 HHPred results for PF3D7_02020700*

| Description | Species | E-value | HH-Hit |
|---|---|---|---|
| Geranylgeranyl pyrophosphate synthase | Arabidopsis thaliana | 5.8e-38 | 3APZ_A |
| Geranylgeranyl pyrophosphate synthase | Corynebacterium glutamicum | 2.2e-37 | 3LMD_A |
| Trans-hexaprenyltransferase | Pseudoalteromonas atlantica | 2.8e-37 | 4JXY_A |
| Hexaprenyl diphosphate transferase | Micrococcus luteus | 3.7e-37 | 3AQB_D |
| Decaprenyl diphosphate synthase | Rhodobacter capsulatus | 3.7e-37 | 3MZV_A |
| Octaprenyl diphosphate synthase | Escherichia coli | 4.4e-37 | 3WJK_B |
| Polyprenyl synthase | Acinetobacter baumannii | 4.4e-37 | 4LOB_A |
| Trans-isoprenyl diphosphate synthase | Caulobacter crescentus | 7.6e-37 | 3OYR_A |
| Geranylgeranyl pyrophosphate synthase | Lactobacillus brevis | 1.1e-36 | 3PKO_A |
| Farnesyl pyrophosphate synthase | Staphylococcus aureus | 1.7e-36 | 5H9D_A |

*excluding hypothetical proteins

**Figure 5- figure supplement 3.** Results of sequence-similarity searches for PF3D7_0202700 using NCBI BLAST and MPI HHpred.

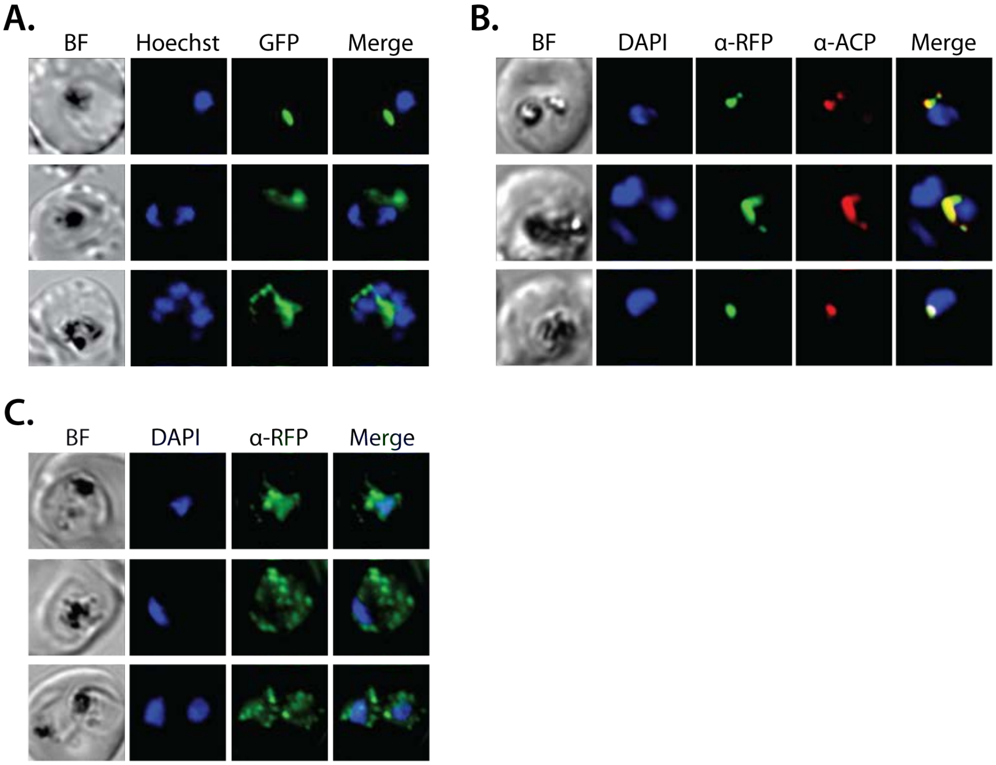


**Figure 5- figure supplement 4.** Additional epifluorescence microscopy images of Dd2 parasite episomally expressing PPS-GFP or PPS-RFP. **(A)** Additional images of live parasite expressing PPS-GFP. Additional immunofluorescence images of Dd2 parasites episomally expressing **(B)** PPS-RFP and stained with anti-apicoplast ACP and an anti-RFP antibody and **(C)** PPS-GFP, cultured for 7 days in 2 μM doxycycline and 200 IPP, and stained with an anti-GFP antibody.

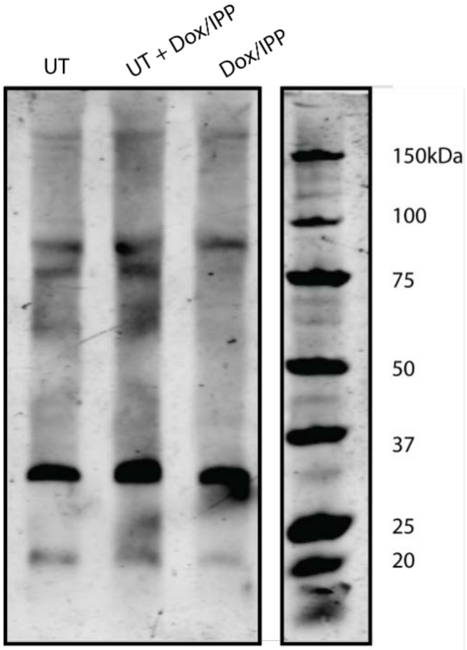


**Figure 5- source data 1.** Uncropped western blot image detecting PPS-RFP expression in parasites.

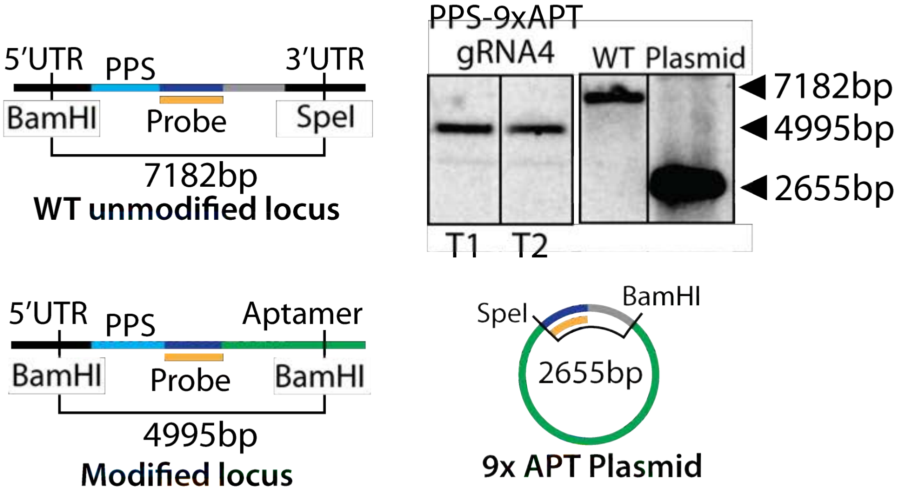


**Figure 6- figure supplement 1.** Scheme for modification of the PPS genomic locus to integrate the aptamer/TetR-DOZI system and Southern blot confirming correct integration.

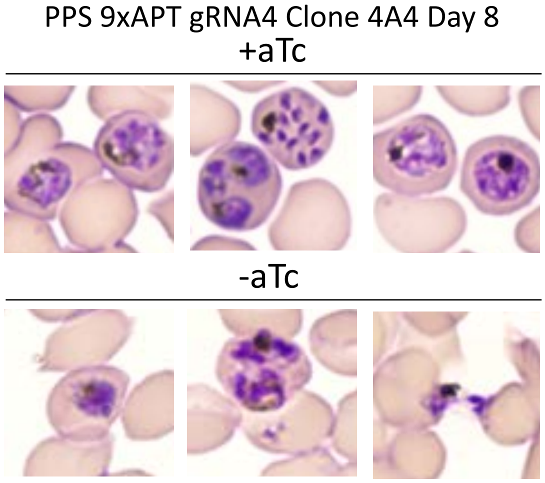


**Figure 6- figure supplement 2.** Blood-smear images of Dd2 parasites tagged at the PPS locus with the aptamer/TetR-DOZI system and grown ±aTc for 8 days.

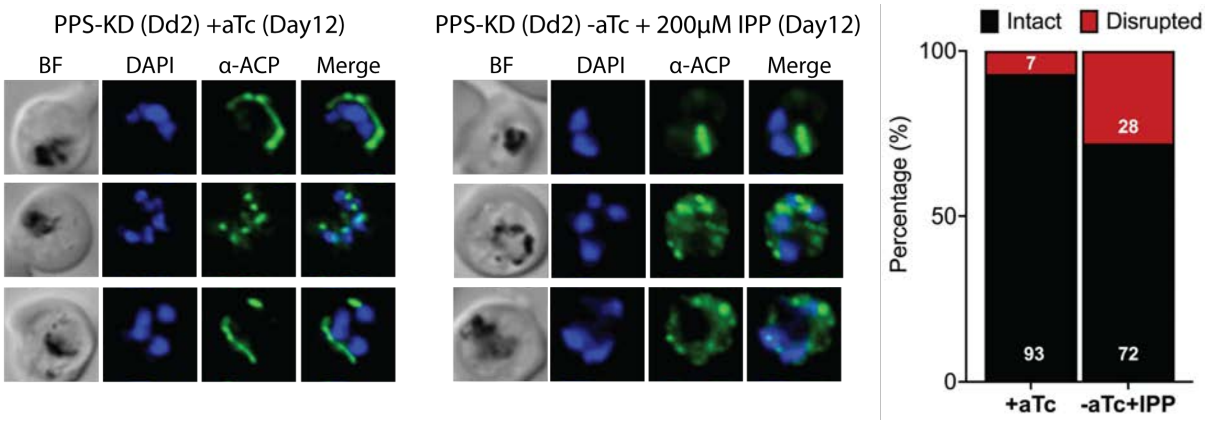


**Figure 7- figure supplement 1.** IFA images and analysis of apicoplast morphology in PPS knockdown parasites grown +aTc or -aTc/+IPP (200 μM) for 12 days. Approximately 30 parasites were imaged in each condition.

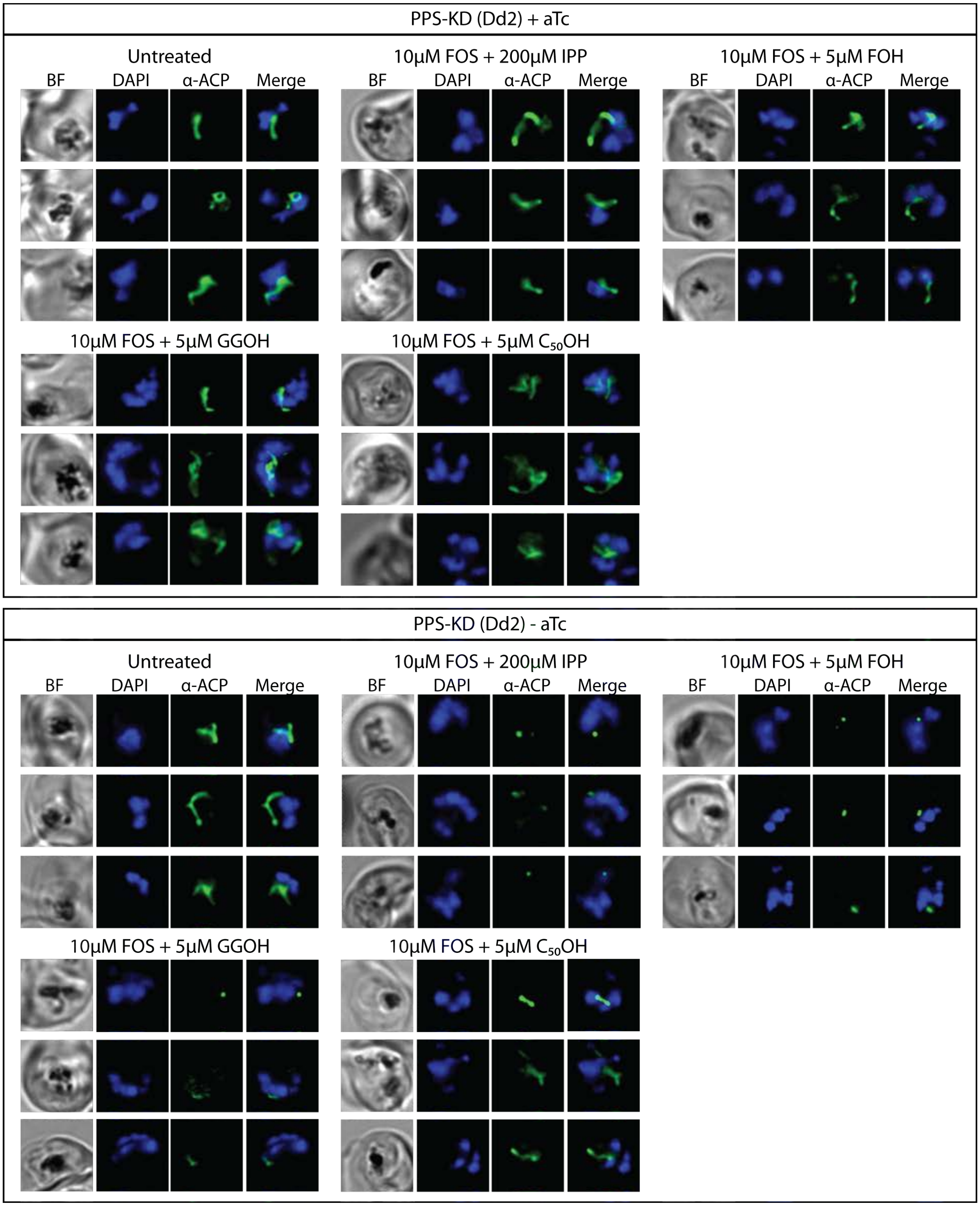


**Figure 7- figure supplement 2.** Additional IFA images of PPS knockdown parasites treated as in Figure 7B.

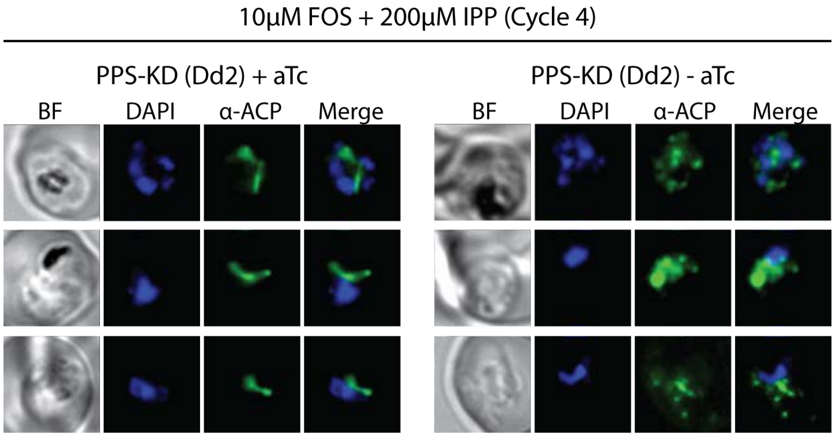


**Figure 7- figure supplement 3.** Additional IFA images of PPS knockdown parasites treated as in Figure 7D.

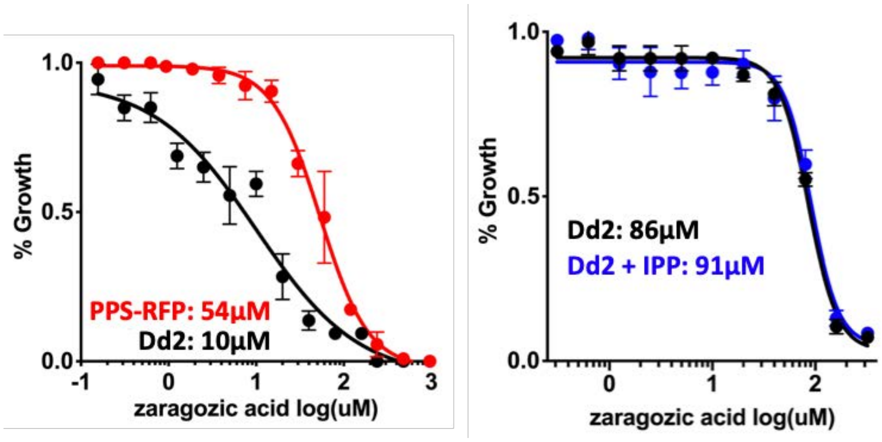


**Figure 8- figure supplement 1.** 48-hour growth inhibition curves for treatment of Dd2 parasites with zaragozic acid without or with episomal expression of PPS-RFP or 200 μM IPP. Parasitemia values were determined for biological duplicate samples at each condition, plotted as the average ±SD, and fit to a four-parameter dose-response curve in GraphPad Prism 9.0.

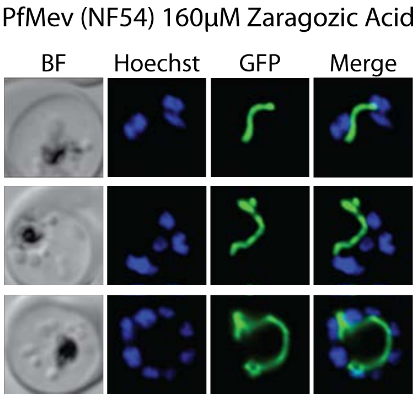


**Figure 8- figure supplement 2.** Epifluorescence microscopy images of D10 parasites treated with 160 μM zaragozic acid as synchronized rings and imaged for $ACP_L$-GFP and Hoescht 36 hours later as multinuclear schizonts.

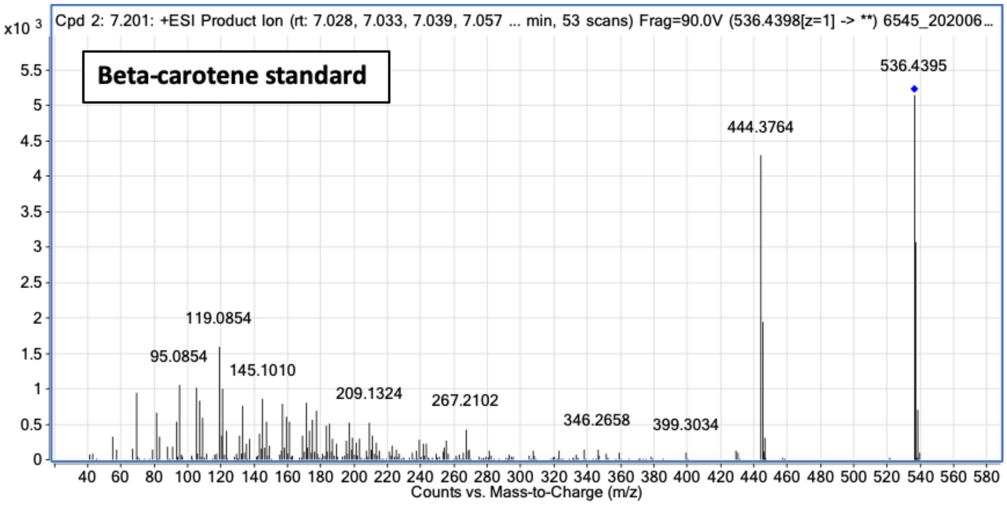


**Figure 8- supplement 3.** Fragment ion spectrum for unlabeled beta-carotene determined by tandem mass spectrometry of β-carotene commercial standard.

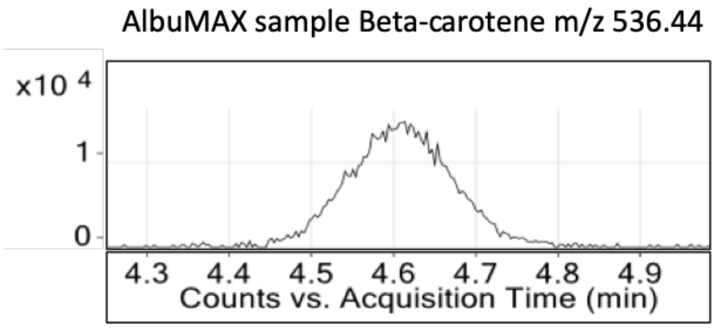


**Figure 8- supplement 4.** Intensity versus retention time plot for liquid chromatography-mass spectrometry determination of unlabeled β-carotene in Albumax I
